# Characterizing metabolism from bulk and single-cell RNA-seq data using METAFlux

**DOI:** 10.1101/2022.05.18.492580

**Authors:** Yuefan Huang, Vakul Mohanty, Merve Dede, May Daher, Li Li, Katayoun Rezvani, Ken Chen

## Abstract

Cells often alter metabolic strategies under nutrient-deprived conditions to support their survival and growth. Characterizing metabolic reprogramming in the TME (Tumor Microenvironment) is of emerging importance in ongoing cancer research and therapy development. Recent developments in mass spectrometry (MS)-based technologies allow simultaneous characterization of metabolic features of tumor, stroma, and immune cells in the TME. However, they only measure a subset of metabolites and cannot provide in situ measurements. Computational methods such as flux balance analysis (FBA) have been developed to estimate metabolic flux from bulk RNA-seq data and have recently been extended to single-cell RNA-seq (scRNA-seq) data. However, it is unclear how reliable the results are, particularly in the context of tissue TME characterization. To investigate this question and fill the analytical gaps, we developed a computational program METAFlux (METAbolic Flux balance analysis), which extends the FBA framework to infer metabolic fluxes from either bulk or single-cell transcriptomic TME data. We benchmarked the prediction accuracy of METAFlux using the exometabolomics data generated on the NCI-60 cell lines and observed significant improvement over existing approaches. We tested METAFlux in bulk RNA-seq data obtained from various tumor types including those in the TCGA. We validated previous knowledge, e.g., lung squamous cell carcinoma (LUSC) has higher glucose uptake than lung adenocarcinoma (LUAD). We also found a novel subset of LUAD samples with unique metabolic profiles and distinct survival outcome. We further examined METAFlux on scRNA-seq data obtained from coculturing tumor cells with CAR-NK cells and observed high consistency between the predicted and the experimental (i.e., Seahorse extracellular) flux measurements. Throughout our investigation, we discovered various modes of metabolic cooperation and competition between various cell-types in TMEs, which could lead to further target discovery and development.

## 1 Introduction

Proper metabolic regulation is essential for the functioning and health of normal cells. However, cancer cells have aberrant genetic alterations, such as amino acid substitutions and copy number alterations, that can perturb cellular metabolism[1, 2]. Thus, cancer cells often exhibit distinct metabolic programs from normal cells. Furthermore, proliferating cancer cells must balance diverged catabolic and anabolic requirements to maintain homeostasis while simultaneously increasing cellular mass [3]. Due to such importance, metabolic dysregulation has been firmly considered a hallmark of cancer[4].

Clinical studies have demonstrated that metabolism is associated with patient outcomes and that specific metabolic phenotypes could present vulnerabilities for cancer treatment[5]. Therefore, understanding how metabolic dysregulation promotes cancer is key to normalizing aberrant metabolism in cancer. However, there are several significant challenges in studying the metabolism of cancer cells in culture. First, it is challenging to mimic a complex metabolic environment, and findings obtained in cell culture may not be effective in clinical settings [3]. Part of the reason is that a tumor microenvironment is a dynamic mixture of malignant and non-malignant cells, which is difficult to mimic in an in-vitro experiment [6]. Also, traditional cell culture nutrient milieu does not resemble human physiological nutrient environment, but nutrient availability in a tumor microenvironment can modulate metabolic dependencies [7, 8]. Over the past two decades, technological innovation in metabolism research has seen tremendous growth. LC/MS (liquid chromatography/mass spectrometry) based metabolomics is a powerful tool for measuring concentration of metabolites and is often the first choice of metabolism experiments [9]. However, the process of assigning peaks into metabolite identity is still low throughput, and it requires lots of time and effort on analyzing the datasets [10].

Moreover, reproducing metabolomics experiments is challenging due to lack of methodology standardization and complexity of experiment[11]. Moreover, metabolomics only provides static metabolic snapshots of cells. Dynamic profiles of metabolite traffic, or velocities of metabolic reactions(fluxes), are critical to understanding mechanisms of cellular metabolic regulation from different angles. Therefore, metabolic flux techniques, for instance, ^13^C metabolic flux analysis (^13^C-MFA) and Seahorse flux assay, are also widely used in metabolic research[10]. ^13^C-metabolic flux analysis (^13^C-MFA) is the current gold standard to measure the intracellular fluxes of central carbon metabolism experimentally. However, it is insufficient to derive large metabolic networks[12]. For extracellular fluxes, Seahorse Extracellular Flux (XF) analyzer has been the industry standard to assess cells’ bioenergetic state. Seahorse metabolic assay can measure OCR (Oxygen consumption rate: an indicator of mitochondrial respiration) and ECAR (extracellular acidification rate: an indicator of glycolysis) of living cells in simultaneously real-time[13]. Even though the Seahorse platform provides valuable insight into the functional status of cells, it only offers two measured fluxes, and interrogating fluxes of other metabolites is not directly available.

With the advent of RNA-Seq and scRNA-seq, we have an opportunity to interrogate the genomic profile of samples with unprecedented high resolution at the system level [14, 15]. Changes in expression of metabolic enzymes, resulting from genetic or downstream pathways alteration or epigenetic modifications [1], provide a valuable snapshot to extend our understanding of tumor metabolism. Recently, transcriptomic analyses have already shown both unique and shared patterns of metabolic reprogramming as well as the metabolic vulnerabilities distinguishing molecular subtypes in various human tumors[16-21].

The most common approach to quantify metabolic pathway activity in transcriptomics data is by summarizing mRNA expression levels of genes in a pathway (i.e., KEGG metabolic pathway gene set) into a single score. Conventional geneset scoring methods can be applied here, for example, ssGSEA[22], AUCell[23], singscore[24], Z-score, etc. However, metabolic network is highly complex, connected, and dynamic, and it may allow cells to overcome inhibition in just a single enzymatic step[25]. Therefore, investigating individual pathways separately without considering the complete map of metabolic circuits is less likely to provide an accurate insight into underlying mechanisms. Moreover, those methods do not produce accurate results with extremely small gene sets. Some of our metabolic reactions contain only one or two genes, so gene set scoring method will be unstable in this scenario. In addition, not all metabolic reactions are associated with enzymes, thus gene set scoring methods are only limited to those with gene association.

Reconstructed Genome-scale metabolic models (GEMs) have been one of the major approaches for systems biology. GEMs encapsulate an organism’s stoichiometric balanced metabolic reactions by gene-protein-reactions (GPR) association[26]. GEMs enable the computational of metabolic response at the system level, which is not possible using metabolic geneset such as KEGG[27]. One of the most common analytic methods used in GEMs is flux balance analysis (FBA), and it predicts the flux values of an entire set of reactions. Flux balance analysis is a well-established constrained optimization method to study such a complicated metabolism by analyzing the flow of metabolites [28-31]. It relies on a stoichiometric representation of the metabolic model to predict the fluxes for reactions while maximizing a cellular objective, subject to constraints, such as the uptake or excretion fluxes observed from the experiment. Furthermore, various extensions of FBA have been successfully applied under different settings[32-37].

Many studies have focused on the integration of FBA and gene expression. They showed such integration could potentially improve the prediction of metabolic fluxes through this low-cost and straightforward fashion[38-49]. One possible solution to connect transcriptome with FBA is using gene expression to define the objective function in FBA. For example, Lee et al. replace the biological objective(i.e., biomass or ATP maintenance) and use the correlation between the fluxes and gene expression score as the objective function[43]. Similarly, iMAT uses gene expression to divide highly expressed and lowly expressed reactions. This method finds the flux distribution that best explains gene expression patterns. It maximizes the number of reactions classified as highly expressed and minimizes the number of reactions classified as lowly expressed[38]. Another way to incorporate gene expression into FBA is to use expression values to define flux bounds in FBA. For example, E-Flux directly uses transformed gene expression values as flux constraints since it assumes the gene expression level determines the flux upper bound[45]. Those studies have demonstrated that the addition of gene expression has improved the prediction of metabolic states [38-42]. However, still lacking is systematic inference of the metabolic reaction activity from mechanistic reaction flux network in human cancer. Current scRNA-seq technology allows powerful characterization of cellular heterogeneity and has unraveled metabolic vulnerability and heterogeneity in many settings[17-19, 50]. Recently, several promising efforts have been made to predict COVID-19 metabolic targets and changes using iMAT on scRNA-seq data [51, 52]. Damiani et al. has proposed single-cell Flux Balance Analysis(scFBA) to obtain single-cell fluxomics. However, usability in large scRNA-seq data was not discussed, and systematic validation in TME was not reported[53].

We developed METAFlux that estimates flux-based metabolic pathway activity from gene expression to address these fundamental challenges. Briefly, we used Human-GEM (consisting of 13082 metabolic reactions, 8378 metabolites) as our underlying metabolic models. Then we derived MRAS (Metabolic Reaction Activity Score) from gene expression using GPR(Gene-protein-reaction)[54] as flux constraints. To account for nutrient differences between cell cultures and patients, we defined lists of nutrient availability profiles for the “cell culture medium” and the “human blood” contexts. Subsequently, we formulated quadratic programming coupled with FBA under given nutrient constraints to calculate the metabolic fluxes for the 13082 reactions. We also extended the framework to calculate single cell cluster level metabolic activity. We combined the different cluster group metabolic network compartments in TME into one network and treated heterogeneous cell clusters as one community, where we can observe different modalities of metabolic organization, such as cooperation and competition for nutrients. Our model produces a higher-resolution activity prediction and will potentially facilitate a more accurate downstream analysis. Notably, our tool has the potential to improve the understanding of aberrant metabolism and serve as a preliminary source to investigate specific metabolic targets.

## 2 Results

### 2.1 Modeling metabolism using transcriptomes

Our underlying metabolic knowledge is extracted from the Human1. Human 1 is a genome-scale metabolic model (GEM) that integrates the Recon, iHSA, and HMR models. It contains 13082 reactions and 8378 metabolites[55]. This model encodes the mechanistic relationships between genes, metabolites, and reactions in a human cell. We choose Human1 because it shows a considerable improvement over other GEMs in terms of stoichiometric consistency, percentages of mass, and charge-balanced reactions[55]. Human1 model contains reactions in 9 compartments (extracellular, peroxisome, mitochondria, cytosol, lysosome, endoplasmic reticulum, golgi apparatus, nucleus, inner mitochondria).

For each sample in a bulk dataset, METAFLux (**Fig. 1a, Methods**) first computes the metabolic reaction activity score (MRAS) for each reaction, which describes the reaction activity as a function of the associated gene expression. Here, we do not consider the reaction kinetic constants and the binding affinity of proteins since robust estimates of all these parameters for genome-scale models are rather difficult[31, 45]. Instead, we use GPR rules, which decode the Boolean logic relationship between genes in a reaction[54], to map the relationship between gene products and then summarize gene expressions into Metabolic Reaction Activity Scores (MRAS) given the predefined relationships. Our approach is adopted from what has been proposed earlier to infer the activity of metabolic reaction from gene expression data [43, 56].

**Figure 1.**
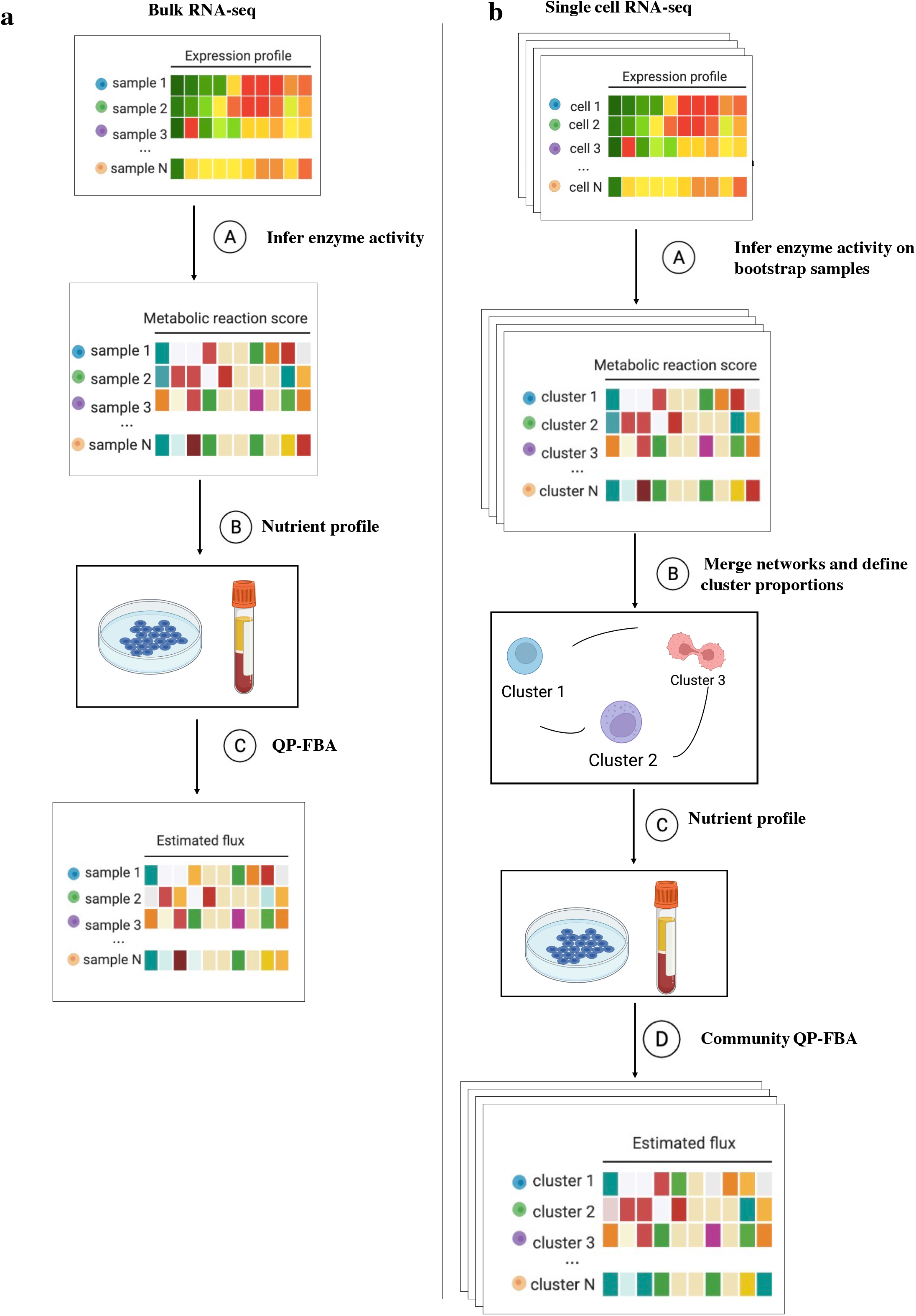
The workflow of METAFlux. **(a)** The workflow of METAFlux in bulk RNA-seq setting. In step A, metabolic reaction activity scores (MRAS) are estimated from RNA-seq data. In step B, a nutrient profile is defined so only certain nutrients can be uptaken. In step C, quadratic programming-based FBA (flux balance analysis) is constructed to estimate metabolic fluxes for each sample. **(b)** The workflow of METAFlux in single-cell RNA-seq setting. In step A, metabolic reaction activity scores (MRAS) are estimated for each stratified bootstrap sampled single-cell dataset. In step B, metabolic networks for different clusters are merged to form one community, and proportions of clusters should be defined during this step. In step C, nutrient profile is defined so only specific metabolites can be uptaken by TME. In step D, community-based quadratic programming FBA is constructed to estimate per cell average metabolic fluxes for each cluster and total average metabolic fluxes for overall TME.

To connect transcriptome and fluxes, a possible solution is to use the MRAS calculated before to define the flux constraints. We use a E-Flux type of approach [45], where the expression levels of the genes associated with a metabolic reaction serves as the maximum possible flux that reaction can carry. The rationale is that, although enzyme activities do not have a high correlation with RNA levels, given a specific level of translational efficiency and assuming there is a limited accumulation of enzymes over a certain time window[45], RNA expression levels can be used as the maximum amount of protein products available and the maximum protein products can then serve as the maximum reaction fluxes.

Subsequently, we need to define a nutrient environment profile, which includes a list of metabolites available for uptake by the reactions (**Methods**). In the constrained optimization step, we hypothesize that tumors proliferate rapidly; thus, the new human biomass pseudo-reaction, which constructs a generic human cell’s nutrient demand and composition, should be optimized [55]. Here we reformulate this idea into convex quadratic programming (QP) to overcome degenerate solutions. Our optimization simultaneously optimizes the biomass objective and minimizes the sum of fluxes’ squares, similar to a previous approach [57](**Methods**).

We also propose a workflow for single cell settings (**Fig. 1b, Methods**). Single-cell RNA-sequencing enables characterization of individual cells to unravel complexity and heterogeneity of tumor microenvironments [58]. Since we believe that cell groups do not work in isolation, modeling the whole tumor microenvironment as one community will account for metabolic interaction between the groups. Here, we do not compute cell-wise flux. Instead, we model fluxes at pseudo-bulk level for the following reasons. First, scRNA-seq data is often highly sparse and noisy. Directly estimating cell-wise MRAS from zero-inflated data can result in many zeros, which challenges downstream modeling. One possible solution seen in the Compass metabolic model, recently developed by Wanger et al., used KNN smoothing to mitigate sparsity and stochasticity [59]. Still, different imputation algorithms can generate different results, an issue beyond the scope of this study. Secondly, merging pseudo-bulk level groups into one network is more computationally efficient than working at single cell level.

To estimate fluxes from scRNA-seq data, we first create stratified bootstrap sampling datasets and compute the pseudo-bulk gene expression profiles based on customized grouping for each bootstrap sample. Next, we calculate MRAS for each pseudo-bulk bootstrap sample. To run METAFlux in single-cell setting, we need to provide cluster (group) proportions with respect to the whole TME. Ideally, these proportions should be retrieved from experiments or calculated from matched bulk data using CIBERSORTx[60]. However, most datasets do not have such information. As a result, directly observed cluster (group) fractions in single-cell data could be used for the purpose, but further studies are warranted to evaluate the findings since those proportions may deviate from the truth due to sparse sampling [61, 62]. We then estimate the group proportions and derive fluxes for each bootstrap using the merged metabolic networks of different groups in TME under constrained optimization (**Methods**).

### 2.2 Benchmarking the performance of METAFlux using experimental data

We used NCI-60 RNA-seq data and publicly available metabolite CORE (<u>co</u>nsumption and <u>re</u>lease rate) data generated on NCI-60 cell lines to benchmark our model performance[63]. However, we only selected 11 cell lines as the experimental ground truth since other cell lines had nutrient depletion that could affect the reliability of flux profiling [55, 64]. There were 26 metabolites fluxes and one biomass flux per cell line. Our model was run based on Ham’s medium composition (**Methods**).

Meanwhile, we also compared METAFlux performance with the state of art method enzyme-constrained model(ecGEMs). These two results can be compared because they were generated based on the same medium composition and both models maximized biomass reaction. ecGEMs is an enzyme-constrained cell line-specific model, which utilizes tINIT to generate cell-specific GEMs and then incorporates GECKO to add enzyme constraints. tINIT is an optimization algorithm that extracts context-specific and connected GEMs based on proteomics and/or transcriptomics datasets[65]. GECKO (enhancement of a Genome-scale model with Enzymatic Constraints and Omics data) extends the flux balance analysis approach (FBA) to enable the integration of enzyme and proteomics data[66].

The overall Spearman correlation of experimental fluxes with predicted fluxes for all the metabolites in the 11 cell lines was 0.76 for METAFlux, and 0.45 for ecGEM (**Fig. 2a**). In terms of consistency across different metabolites, METAFlux generally achieved better spearman correlation with ground truth than ecGEM across metabolites (**Fig. 2b)**. For metabolites like L-carnitine and pyruvate, ecGEM predicted those fluxes to be 0. Therefore, the correlation of these two metabolites to background truth was not calculated. To further examine the accuracy of our flux prediction, we categorized metabolic fluxes to ‘no flux’ (flux equals to 0), ‘uptake’ (flux smaller than 0) and ‘excrete’ (flux greater than 0) because correlation metrics only show whether there is a relationship between pairs of fluxes. However, it omits the sign of fluxes which represents the directional uptake or excretion behavior. After categorization, we evaluated the predicted direction accuracy for each metabolite between two models **(Fig. 2c)**.

**Figure 2.**
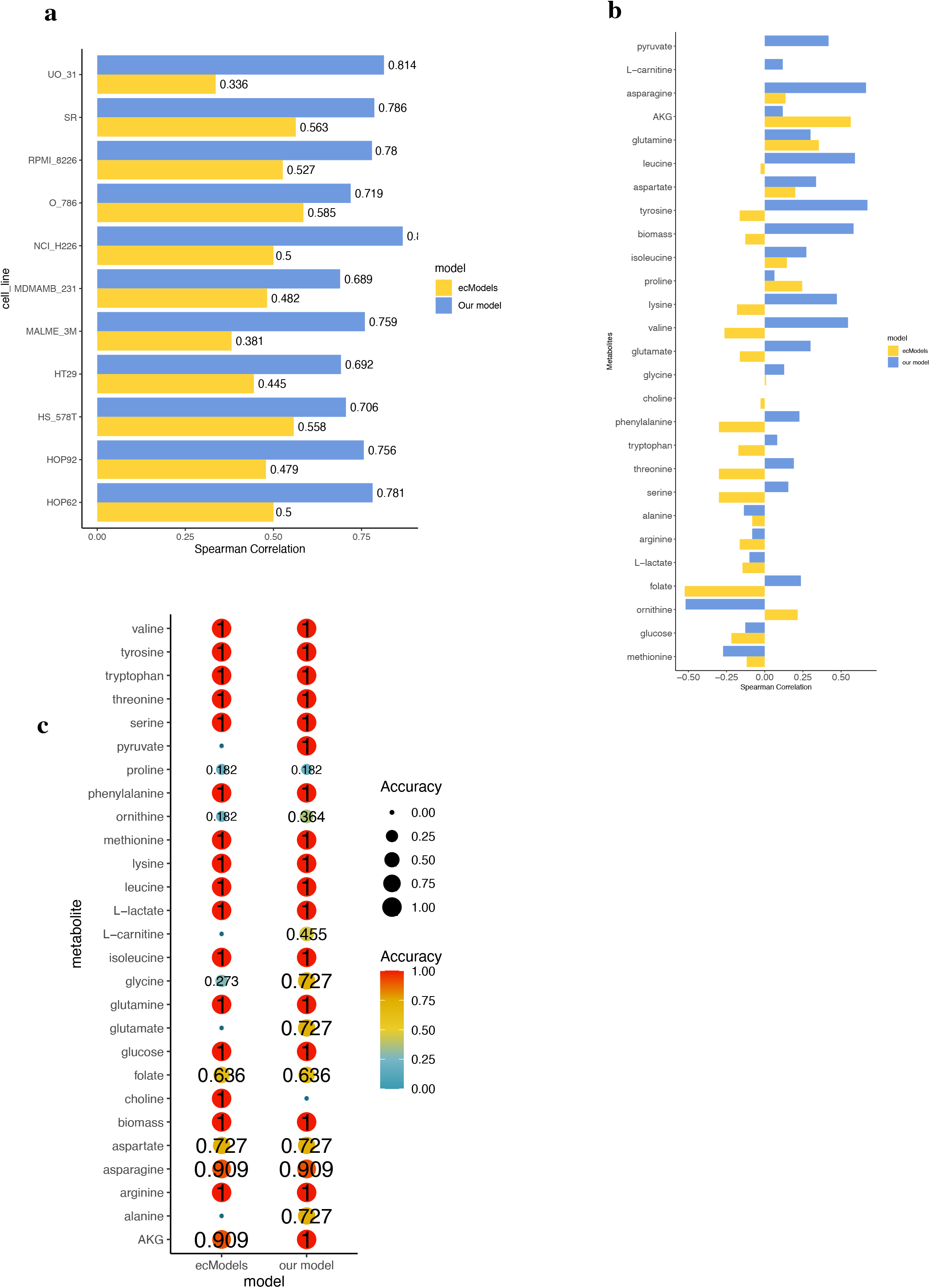
Benchmark results of METAFlux on NCI-60 cell lines and performance comparison with ecModel. **(a)** Spearman correlation bar plot across each cell line for METAFlux and ecModel. The spearman correlations between predicted fluxes and experimental fluxes were calculated for 11 cell lines. **(b)** Spearman correlation bar plot across each metabolite for METAFlux and ecModel. The spearman correlations between predicted fluxes and experimental fluxes were calculated for 26 metabolites and one biomass reaction. (**c**) Uptake and secretion direction accuracies of 26 metabolites and biomass reaction for METAFlux and ecModel. The accuracies were defined by the ratio of the number of direction-aligned fluxes to the total number of fluxes for each metabolite.

Except for ‘choline,’ METAFlux achieved better accuracy for seven metabolites and the same accuracy with ecGEM for the other 19 metabolites. Taken together, these results suggest that METAFlux outperformed ecGEM on predicting metabolic fluxes from RNA-seq data.

### 2.3 Biological meaningful medium resolved better prediction

To estimate the effect of medium constraints on model performance, we first did not impose medium constraints in METAFlux, meaning that we freely allowed the uptake or excretion of all exchange metabolites with no rate restriction. Without any medium constraints, METAFlux only had a spearman correlation of 0.15. Similarly, both spearman correlations across metabolites and cell lines dropped significantly **(Supplementary Figure 1a, b)**. The result showed that the prediction performance for models without medium constraints declined considerably compared with the original medium composition.

To further explore the medium effect on performance, we tested our model on random generated medium. We compared the results obtained from our biological meaningful medium with same-sized random mediums. The original hypothesized hams medium contains 44 metabolites, so we randomly selected 44 metabolites from the total 1648 exchange metabolites. We then allowed our model to uptake or excrete those 44 metabolites without rate restriction while only allowing the rest of 1604 metabolites to excrete with no rate restriction. We repeated this simulation process *N* = 500 times. For each simulation *i*, we obtained the overall spearman correlation *ρ*_*i*_ and directionality accuracy *acc*_*i*_. To calculate the p-value, we counted the number of measurements greater than our original biological meaningful statistics *ρ*_*o*_ and *acc*_*o*_ and divided this number by 500. The P-values for both overall spearman accuracy and direction accuracy were zero, indicating that none of the results generated by the random medium were superior to our original biological medium **(Supplementary Figure 1c, d)**. These results indicate that METAFlux has achieved biologically meaningful modeling in *in vitro* cell-line culturing experiments and can potentially be applied in broader settings.

### 2.4 METAFlux identified a novel metabolic cluster in TCGA lung cancer

We applied METAFlux on TCGA LUAD (lung adenocarcinoma) and TCGA LUSC (lung squamous cell carcinoma) with human blood profile as medium constraints (**Methods**). A recent study utilizing TCIA (The Cancer Imaging Archive) database to quantify tumor glucose uptake using ^18^F-FGD PET-CT observed a significant higher glucose uptake in LUSC than in LUAD. To validate METAFlux, we examined the glucose uptake flux estimated by METAFlux from the LUSC and the LUAD RNA-seq data. The METAFlux prediction results indicated that LUSC tumors had a higher glucose uptake than LUAD, which is consistent with the ^18^F-FGD PET-CT scan results **(Fig. 3a)**. In addition, prior research has also suggested ^18^F-FGD was closely correlated with the proliferation index[67]. Consistent with this, we found that the glucose uptake influx from METAFlux was highly correlated with estimated proliferation signature scores with a spearman correlation of 0.65 **(Supplementary Figure 2a)**. The proliferation scores were calculated by ssGSEA using derived gene sets and were directly available from earlier study [68].

**Figure 3.**
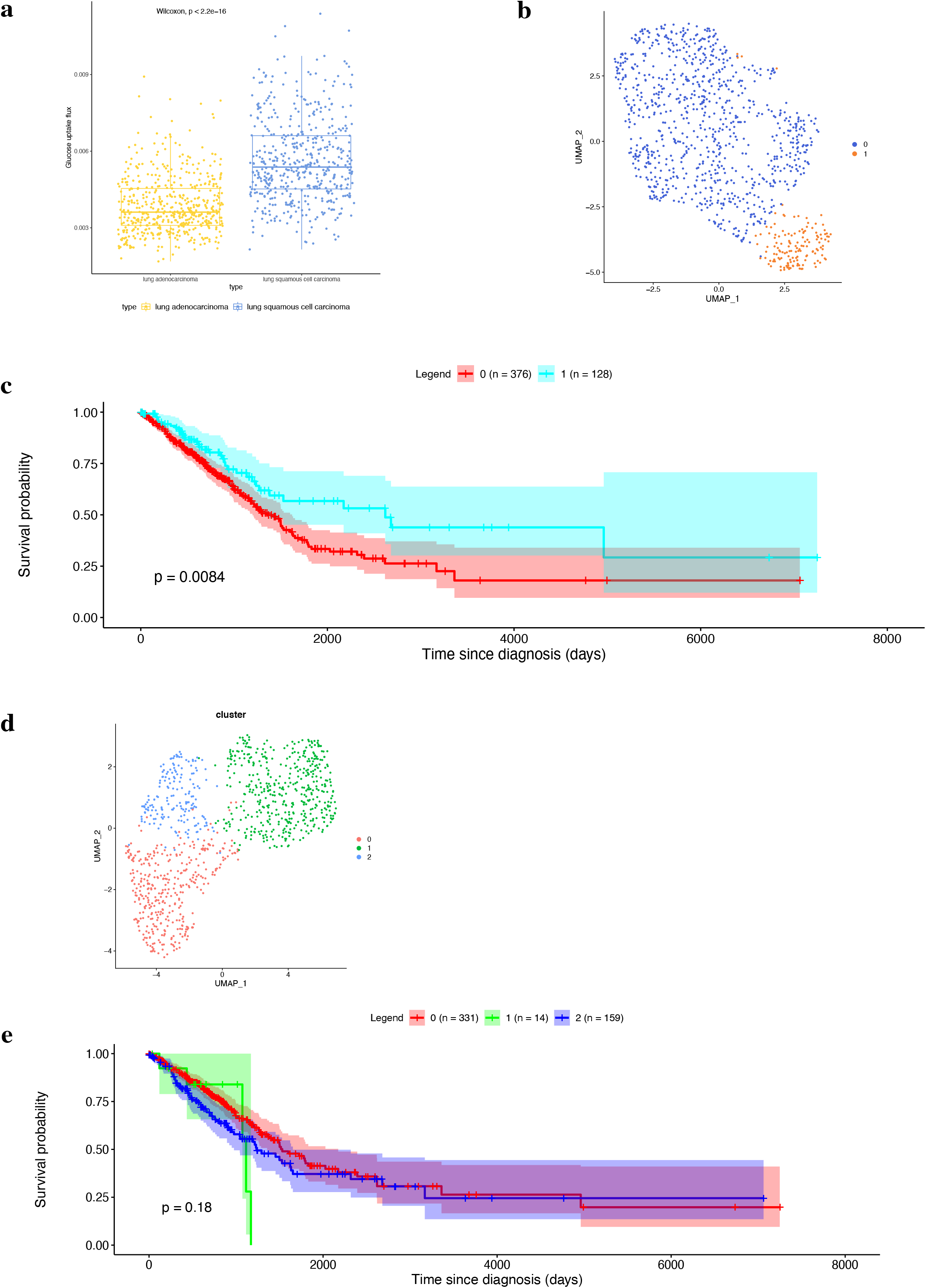
METAFlux application on TCGA LUAD and LUSC datasets. **(a)** Boxplot of glucose uptake activity in LUAD and LUSC. **(b)** UMAP of TCGA LUAD and LUSC samples using predicted metabolic fluxes. Two clusters were identified within those samples. Cluster 1 was enriched with LUAD tumors, while cluster 0 contained both LUAD and LUSC tumors. **(c)** Overall Kaplan-Meier survival curves for clusters 0 and 1 in TCGA LUAD. Clusters were generated using metabolic fluxes. **(d)** UMAP of TCGA LUAD and LUSC samples using metabolic gene expression. **(e)** Overall Kaplan-Meier survival curves for clusters in TCGA LUAD. Clusters were generated using metabolic gene expression.

Clustering the metabolic fluxes revealed 2 clusters of samples for LUAD and LUSC **(Fig. 3b)**. Cluster 1 was primarily made up of LUAD tumors, while cluster 0 was a mixture of LUAD and LUSC tumors **(Supplementary Figure 2b)**. The LUAD tumors in cluster 0 are called ‘LUSC-like LUAD,’ because this subset of LUAD tumors exhibited a similar metabolic phenotype with LUSC. The LUAD tumors in cluster 1 (LUAD1) had significantly lower glucose uptake than ‘LUSC-like LUAD” (*P* values<2.26× 10^−16^), implying LUSC-like LUAD tumors are more metabolically active. Survival analysis revealed that LUAD1 tumors had significantly better survival outcomes (P value < 0.0084) than ‘LUSC-like LUAD’ **(Fig. 3c)**. Moreover, such clustering membership cannot be found when performing clustering using the corresponding gene expression data **(Fig. 3d)** and **Supplementary Figure 2c, d)**. There was no significant difference in patient survival among the resulting clusters in LUAD tumors **(Fig. 3e)**. Results demonstrated that utilizing metabolic fluxes generated by METAFlux was able to discover novel tumor subtypes, which could not be found otherwise by gene expression.

### 2.5 METAFlux identified a longitudinal metabolic competition trend between Tumor and NK cells

In single cell settings, we first examined the scRNA-seq data generated from an *in vivo* coculture experiment assessing the killing of engineered CAR-NK cells on Raji cells, a non-curative CD19+ lymphoma cell-line model. Data included three products: CAR19-NK cells armed with IL-15, CAR19-NK cells lacking IL-15, and non-transduced NK cells, in addition to the tumor cells. In total, data from four time points were collected: day 7, day 14, day 21, and day28. The extracellular acidification rate (ECAR) and oxygen consumption rate (OCR) data were measured by Agilent Seahorse XFe96 Analyzer, and the measurements were taken 2 hours after cocultured NK and Raji cells. METAFlux single cell framework requires cell type proportion as an input parameters, here we simply calculated percentage for each cell type using single cell RNA-seq data.

We sought to compare our METAFlux results with the Seahorse assay. We only utilized single-cell RNA-seq data from day 7 for our benchmark comparison since day 7 profile would be the closest to the state when Seahorse assay was measured. We used oxygen efflux as the surrogate for OCR. Similar to a previous study, we used proton efflux as the surrogate for ECAR[69]. We set the bootstrap number to 100 for each product. The Seahorse assay indicated that CAR19/IL15 NK cells had the highest OCR and ECAR, followed by CAR19 NK cells, and NT-NK cells showed the lowest OCR and ECAR among the three products. Our predicted OCR and ECAR findings showed a consistent trend with Seahorse Assay that CAR19/IL15 NK cells were the most metabolically active, while NT-NK cells were the least metabolically active **(Fig. 4a, b** and **Supplementary Figure 3a-b)**.

**Figure 4.**
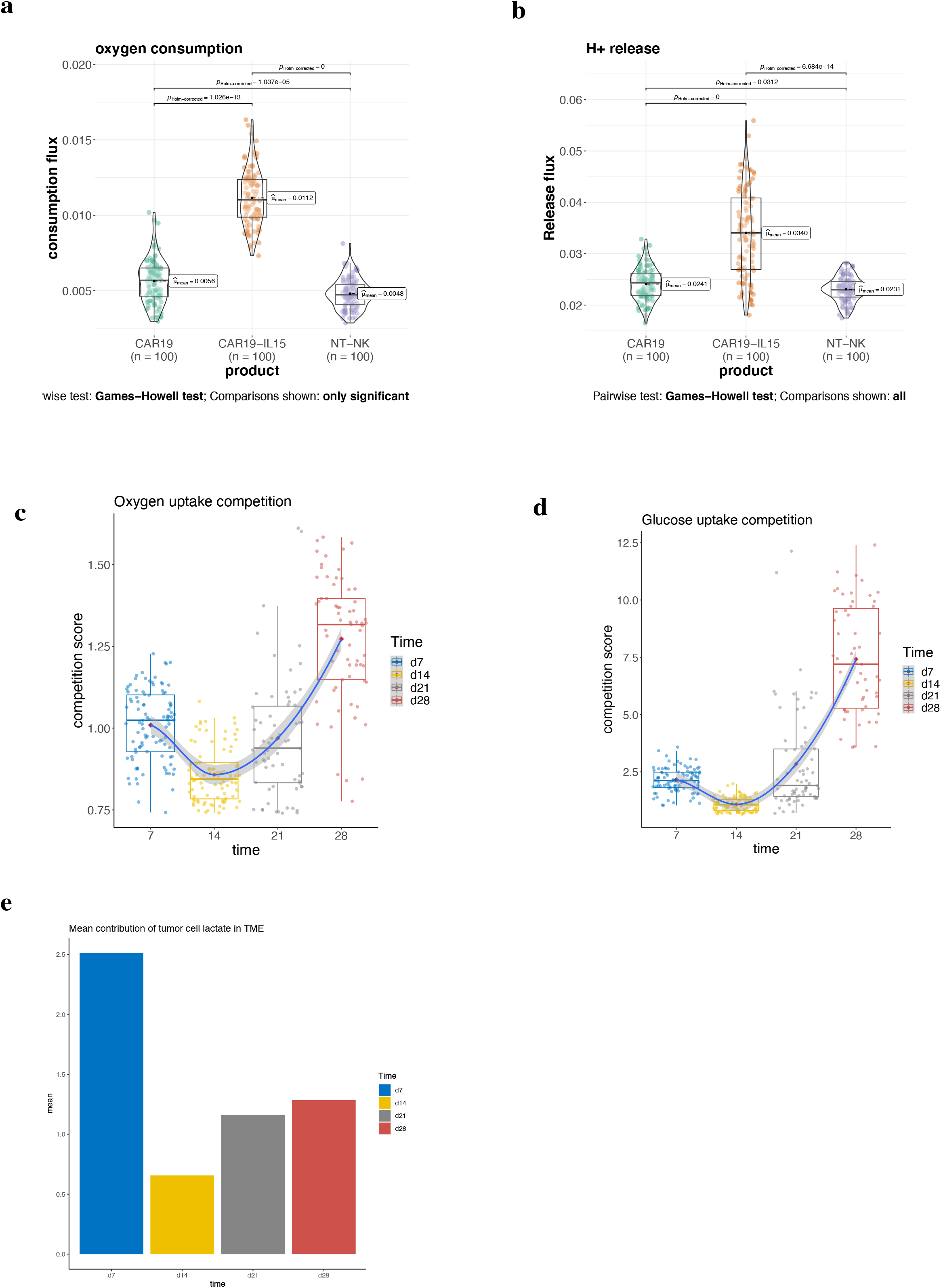
CAR-NK single-cell RNA-seq METAFlux analysis. **(a)** Violin plot of oxygen consumption flux for the day 7 product of CAR19-IL15, CAR19, NT-NK. Each group includes n=100 bootstrap samples. **(b)** Violin plot of H+ release flux for the day 7 product of CAR19-IL15, CAR19, NT-NK. Each group includes n=100 bootstrap samples. (c-d) Competition score of (c) glucose and (b)oxygen for average cancer cell against average NK cell in TME from day 7 to day 28. **(e)** Barplot of tumor cells contribution of lactate release in TME.

Next, we aimed to analyze the metabolic competition in CAR19/IL15 over time. Our ongoing work demonstrated tumors recurred after day 14, and NK cells experienced metabolic dysfunction and decreased anti-tumor activity over time with tumor recurrence. We set the bootstrap number for each time point to be 100 and used per cell competition score to quantify the metabolic competition between tumor and NK cells. We defined tumor and NK nutrient competition score as the ratio of the per cell nutrient uptake flux in tumor to the per cell nutrient uptake flux in NK. We found that the metabolic competition for oxygen and glucose decreased from day 7 to day 14 but ramped up after day 14 and reached its peak on day 28**(Fig. 4c, d)**. Moreover, several amino acids showed similar trends **(Supplementary Figure 3c-f)**. Consistent with prior work, METAFlux results suggested tumor cells eventually outcompeted NK cells for nutrients and NK cells became less metabolically fit over time[70].

We also observed that tumor cells were the major contributor to the total lactate production in the TME on day 21 and day 28, and there was an increasing trend of lactate production by all tumor cells over time **(Fig. 4e** and **Supplementary Figure 3g)**. On average cell level, NK cells showed an elevated lactate release over time **(Supplementary Figure 3h)**. As a result, the tumor microenvironment became more and more acidic, which suppressed the NK function and decreased the cytotoxic activity of NK cells, leading to tumor recurrence[71]. Our findings could be potentially relevant to understanding the associated mechanism with tumor resistance and relapse. Taken together, METAFlux can capture the longitudinal competition mechanism between NK and tumor cells and pinpoint the nutrients they compete for.

### 2.6 METAFlux identified different modalities of metabolism

Our TCGA sample analysis demonstrated that glucose uptake was significantly higher in LUSC than in LUAD at bulk level. However, glucose uptake comparison on bulk level did not show the distinct metabolic programs for different cell types in TME. To this end, we then sought to identify metabolic heterogeneity for different cell types using METAFlux in a community-based setting.

We applied METAFlux on bulk sorted RNA profiling data from primary lung cancer patients directly acquired from the operating room[72]. The data was sorted into immune cells (CD45+EPCAM−), endothelial cells (CD31+CD45−EPCAM−), tumor cells (EPCAM+CD45−CD31−), and fibroblasts (CD10+EPCAM−CD45−CD31−). After data processing, we have 15 ADENO(lung adenocarcinoma) and 9 SCC (lung squamous cell carcinoma) samples with complete RNA-seq profiles for all four cell types. We used the cell type proportions from the original study as our input parameters and those cell type proportions were calculated by CIBERSORT using matched bulk data[72].

We next compared the nutrient uptake between cancer, immune cell, fibroblast, and endothelial cells between SCC and ADENO. Since the sample size was small, the P value was not significant. Consistent with the findings from the bulk data, there was a trend toward higher glucose uptake from whole TME in SCC than in ADENO (*P*=0.058,95% CI of effect size [-1.8149, - 0.1191]) **(Supplementary Figure 4a)**. The cell type specific nutrient uptake indicated a weaker increase in glucose uptake from average cancer cells in SCC than in ADENO (*P*=0.222, 95% CI of effect size[-1.3435,0.2831]) **(Fig. 5a)**. The glucose uptake per cell in endothelial and fibroblast cells was similar between SCC and ADENO (endothelial *P*=0.762, 95% CI of effect size [-0.9300,0.6677], fibroblasts *P*=0.803,95% CI of effect size [-0.9037,0.6934]) **(Fig. 5a)**. However, the most striking difference came from immune cells. The immune cells in SCC had the highest per cell glucose uptake in TME, while the immune cells in ADENO had the lowest capacity to uptake glucose per cell among TME (*P*=0.045, 95% CI of effect size[-1.8626, -0.1579]) **(Fig. 5a)**. This finding suggested that immune cells in SCC were not deprived of glucose.

**Figure 5.**
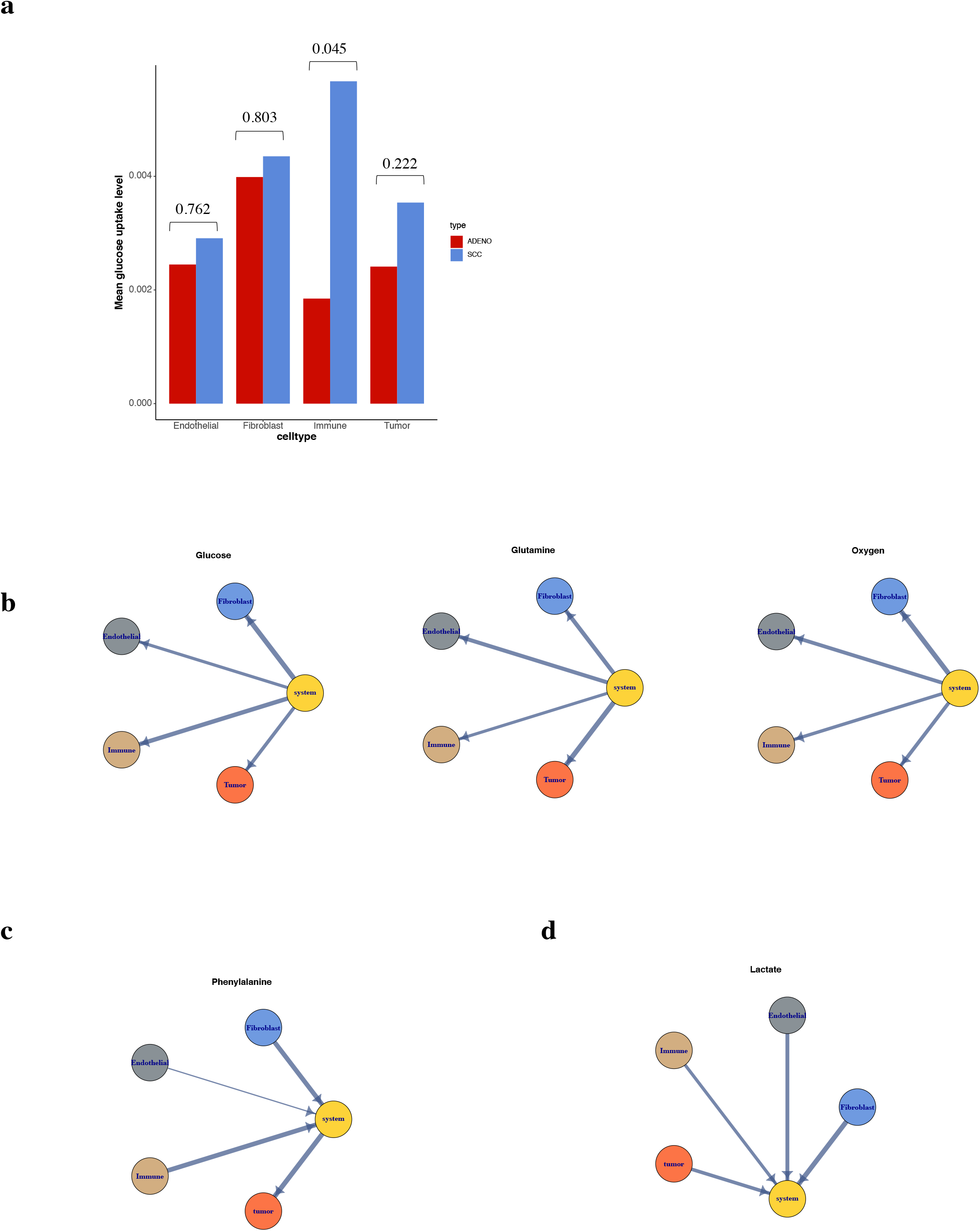
Different modalities of metabolic mechanisms identified by METAFlux. **(a)** Bar plot of per cell average cell type-specific glucose uptake fluxes for ADENO and SCC. (**b-d**) Graph-based representation of 3 different metabolic modalities in ADENO and SCC. Each node represents one cell type (Immune, endothelial, tumor, or fibroblast) or system (shared TME). The edge width represents the absolute magnitude of flux. The arrow shows the direction of flux. Arrow coming from cells to system means the nutrient of interest is released to system. Arrow coming from system to cells means the nutrient of interest is absorbed from system. **(b)** Metabolic competition mode in ADENO and SCC. Immune, endothelial, tumor and fibroblast cells compete for glucose, glutamine, and oxygen uptake in TME (represented by the system). **(C)** Metabolic cooperation mode in ADENO and SCC. Immune, endothelial, and fibroblast cells release phenylalanine in TME (represented by the system) and tumor cells uptake phenylalanine to favor their growth. (d) Parallel producer mode in ADENO and SCC. Lactate released by all cell types to TME (system).

In addition to per cell metabolic profile for each cell type, we were also interested in total nutrient consumption for a particular cell type. To calculate the total consumption for a certain nutrient in a particular cell type, we multiplied the per cell nutrient uptake flux by its cell type abundance. Since the tumor has the largest abundance, the nutrients secretion or uptake can be greatly skewed by its mass. Cancer cells account for 56.55% of glucose uptake in ADENO and 49.28% of glucose uptake in SCC. Immune cells account for 18.82% of glucose uptake in ADENO and 34.31% of glucose in SCC**(Supplementary Figure 4b)**. For glutamine, cancer cells account for 68.25% of glutamine uptake in ADENO and 62.32% of glutamine uptake in SCC. Immune cells account for 15.79% of glutamine uptake in ADENO and 19.33 % of glutamine in SCC **(Supplementary Figure 4b)**. METAFlux showed that heterogeneous cell types had different affinities for nutrient and revealed distinct metabolic phenotypes within TME.

TME is a highly complex mixture, and TME components either form metabolic antagonism or symbiosis when uptaking nutrients[73]. When one or more cell types benefit from the metabolites produced by other cell types, we define the interaction as a metabolic cooperation program. When all cell types compete for limited resources in TME, we define it as a competition metabolic program. A competition between TME components occurs when nutrient demands in TME are high.

We captured different interaction mechanisms among TME in both ADENO and SCC **(Fig. 5b-d)**. We identified the competition mode where different cell types compete for nutrients like glucose, glutamine, oxygen, and other amino acids **(Fig. 5b** and **Supplementary Tables 1)**. In contrast, we have also identified a cooperation mechanism where one or more cell types utilized the nutrients (e.g., Phenylalanine) produced by other TME components to favor their growth **(Fig. 5c** and **Supplementary Tables 1)**. Besides cooperation and competition, we also found a ‘parallel producer’ modality where all TME compartments released a certain nutrient (e.g., lactate) (**Fig. 5d)**.

## 3 Discussion

In this study, we developed METAFlux, a computational framework that opens up the application of metabolic flux analysis to transcriptomic datasets. First, we converted RNA-seq data to metabolic reaction activity score and use constrained optimization to maximize biomass under the steady-state assumption to infer fluxes. We then extended the method to a single-cell RNA-seq setting. For single cell settings, we first merged different compartments in TME into one network and calculate cluster level metabolic activity. Then, we treated heterogeneous cell clusters in TME as one community and used constrained optimization to maximize biomass for the entire community and infer metabolic fluxes.

METAFlux calculates metabolic fluxes at cluster level under single cell settings, because we observed that RNA-count dropout could affect flux prediction accuracy. Creating a pseudo-bulk sample can substantially alleviate the negative impact of dropout effect and save computational cost. For datasets with high dropout rates, it could be beneficial to applying gene expression imputation before applying METAFlux.

The metabolic growth highly relies on the medium; however, we did not distinguish different levels of nutrient concentrations in the medium of METAFlux, and we only utilized the binary condition (presence vs. absence) of nutrients in medium as a constraint. This approach can be helpful when detailed experimental parameters are not available. However, there could be situations where the concentrations or fluxes of certain nutrients are significantly different for each sample, for example, different levels of hypoxia experienced by samples. Thus, more realistic profiling of the medium should increase the prediction accuracy in that case.

Therefore, our future development could focus on the extension to enable quantitative integration of experimental nutrient concentration or measured metabolic fluxes.

METAFlux in bulk RNA-seq setting achieved great benchmarking results using NCI 60 experimental data as ground truth. It correctly predicted metabolic phenotypes in single cell CAR-NK dataset, compared against Seahorse assays. We also successfully applied METAFlux on various patient samples for hypothesis generation. Our tool provides a cost-effective approach for discovering novel metabolic features in high-dimensional datasets. We have shown our method can generate consistent results with experimental data and identify meaningful and novel features. Furthermore, our tool allows for the connection between transcriptomic data and system-level FBA metabolic analysis and a better understanding of plastic and complex metabolic network.

## 4 Methods

### 4.1 Underlying genome-scale metabolic model (GEM)

### 4.2 Inference of metabolic reaction activity score (MRAS) from transcriptomic data

In GPR, the **AND operator** joins the genes encoding for different subunits of the same enzyme, and the **OR operator** joins the genes encoding for isoenzymes[74]. For a reaction catalyzed by an enzyme complex, all the subunits need to be expressed to catalyze a reaction, and the lowest expressed unit will be the rate-limiting step for this complex. Therefore, the metabolic activity of such an enzyme complex will be the lowest expression value among all genes associated with this enzyme complex. For a reaction catalyzed by isoenzymes, all the isozymes contribute additively to this reaction[56]. Thus, metabolic activity will be the sum of all expressions of isoenzyme genes. Some genes are involved in multiple reactions (e.g., promiscuous enzyme), and we hypothesize that there may be enzyme resource competition may exist between reactions. We adjust for the enzyme promiscuity by dividing the expression value of a gene by the number of reactions the gene has participated in. A similar approach has been seen in[75]. The steps of deriving MRAS are the following:

Let *w*_*i*_ be the number of reactions *Enzyme*_*i*_ participate and

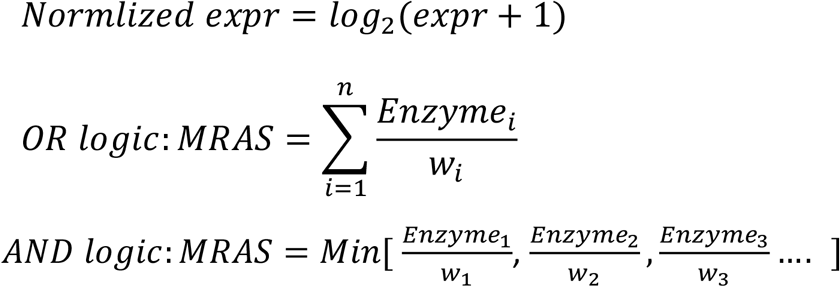

#### 4.2.1 Reaction flux constraints setup

We set normalized MRAS as the flux upper bound to their corresponding metabolic reactions. The lower bound of reaction flux is set to be 0 if the reaction is non-reversible and (−*normalized MRAS*) if the reaction is reversible. The flux is loosely constrained when MRAS is high, so there is more bandwidth of reaction flux. On the other hand, the flux is strictly constrained when MRAS is low, so the bandwidth of flux is much narrower. Our model consists of 13,082 metabolic reactions in total, and 8,033 reactions (8,033 R1 reactions) are associated with enzymes. For the rest of the reactions that are not associated with any enzymes (5,049 R2 reactions), those reactions are not constrained by gene expression, so we set the upper bound as one.

The constraints of flux are as follows:

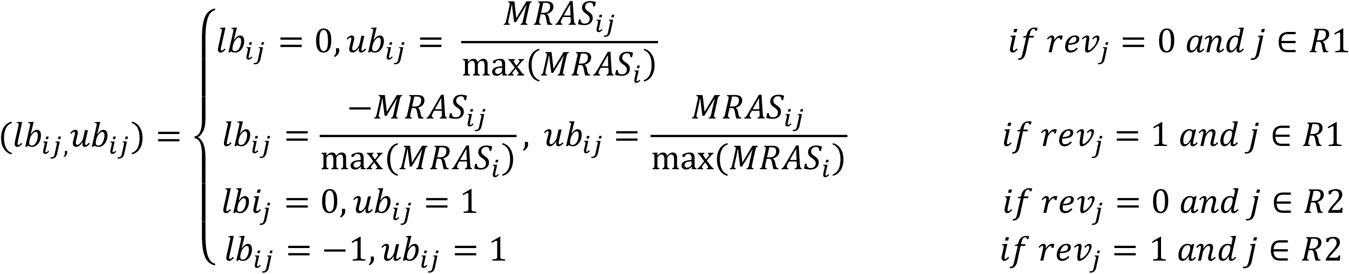

Where *lb*_*ij*_ and *ub*_*ij*_are the input flux bounds for *jth* reaction in *ith* samples, and *rev* stands for reversibility of a reaction. If *rev* = 0, the reaction is non-reversible; if *rev* = 1, the reaction is reversible.

### 4.3 Optimization modeling

#### 4.3.1 Stoichiometric representation of metabolic reactions

A chemical reaction is a basic unit in metabolic pathways, and stoichiometry can represent the quantitative relationship between products and reactants in a reaction. The stoichiometric coefficients of the reactions that constitute the network populate the stoichiometric matrix **S**. Here, Stoichiometric matrix is an 8378 **M** by 13082 **N** sparse matrix, where **M** equals the number of metabolites in different compartments, and **N** equals the total number of metabolic reactions. The negative coefficient refers to the number of moles of metabolites are consumed in a particular reaction. The positive coefficient means how many moles of the metabolites are produced in a specific reaction. At the same time, zero implies this metabolite does not participate in a specific reaction.

#### 4.3.2 Defining nutrient availability profile for cell culture and patient samples

We use the ham’s medium as the growth medium in cell line models [55]. It contains 44 metabolites, and the uptake or excretion rates of these 44 metabolites are not limited, meaning that cells may uptake or excrete these metabolites without limit. For the remaining metabolites in the model, we do not allow cells to uptake from the medium, but cells can excrete those metabolites into the medium. For tissue samples from patients, it is necessary to define a more physiologically relevant environment as the traditional synthetic medium does not mimic human blood. Jason et al. developed a human plasma-like medium [HPLM] to better capture the composition of human blood, and we derive a list of 64 metabolites in human blood based on their profiling[7].

#### 4.3.3 Quadratic programming

Traditional FBA by linear programming (LP) gives a unique optimal objective value. However, the solution to FBA from LP is most likely degenerate, meaning the solutions are not unique, and different solvers will likely return different vectors. Usually, the flux variability analysis will be used afterward to calculate the range of fluxes that achieves the optimal objective[76].

Another approach, pFBA or Parsimonious enzyme usage FBA, was proposed earlier [32]. pFBA assumes there is a selection for an organism to minimize the total amount of necessary enzymes to achieve optimal growth. pFBA first computes the optimal growth rate and then minimizes the sum of reaction fluxes under the optimal solution. Here we reformulate this idea into convex quadratic programming (QP).

We define single-sample unsolved metabolic fluxes by vector **v** with the length of 13082. The dot product of matrix **S** and a vector of unknown fluxes **v** approximation to 0 represents steady-state assumption where essentially the metabolite concentrations are held constant 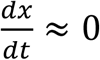, where **x** stands for the concentration of all metabolites.

The framework is implemented using OSQP solver[77] and formulated as the following optimization problem:

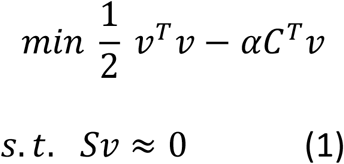

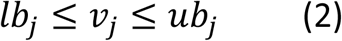

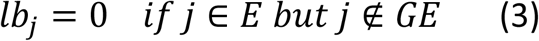

Where *C* is a vector of zeros with a one at the position of our designated biomass reaction, and constraint (3) is the growth medium constraints. *a* is set to 10000, the same order of magnitude with respect to fluxes used in Fit methods[57]. *j* stands for the *jth* reaction. We define a growth medium *G* and all exchange reactions *E* and *GE* as exchange reactions relevant to *G*.

### 4.4 Community-based flux estimation in single-cell data

Given a scRNA-seq dataset, we assign a group (cluster or cell type) label to each cell. We first perform stratified bootstrapping, which means we sampled with replacement with respect to each group.

Step 1: Bootstrap sample generation. Let group be *g* = 1, …. *n*, and *B*_*ig*_ be the *ith* bootstrap sample for *gth* group. For each bootstrap iteration *i*, we combine *B*_*i*1_,*B*_*i2*_… for all groups to form resampled data. Each generated bootstrap data will be the same size and have the same group proportion as the original data.

Step 2: Mean calculation. For each bootstrap sample, we calculate the mean gene expression vector for each group.

Step 3: Define group fraction parameter *P. P* is a fraction matrix defined as:

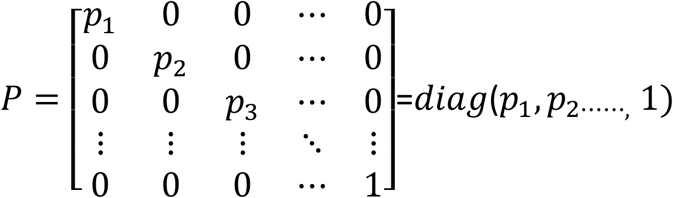

*Pi* stands for the percentage of cell group *i*, and we constrain that 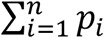 should be one.

We require group fractions as our input. Group fractions indicate the proportions of groups of interest with respect to the whole sample.

Step 4: Merging metabolic networks. Let 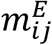 be the metabolites associated with exchange reactions in group *i*. To merge multiple metabolic networks, we need to create a “TME metabolite reservoir” for different cell groups to interact. We define *rm*_*ij*_ as the reservoir metabolite *j* in group *i*

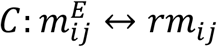

This representation allows different partitions of TME to share the same resources.

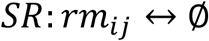

This representation ensures the model is an open system and allows reservoir metabolites to be exchanged with the external environment. The final size of the merged stoichiometric matrix is *(N* × *8378 +* 16*48*) × (1*3*0*82* × *N +* 16*48*), where *N* stands for the number for groups we defined. A specific construct of the merged stoichiometric matrix for three groups is shown below:

|  | Group 1<br>reactions | Group 2<br>reactions | Group 3<br>reactions | Shared<br>metabolic<br>reactions |
| --- | --- | --- | --- | --- |
| Shared<br>metabolites | $C_i$ | $C_j$ | $C_k$ | $SR$ |
| Group 1<br>Metabolites | $S_i$ | **0** | **0** | **0** |
| Group 2<br>Metabolites | **0** | $S_j$ | **0** | **0** |
| Group 3<br>Metabolites | **0** | **0** | $S_k$ | **0** |

Our model aims to maximize the entire community’s biomass while minimizing the sum square of overall fluxes. The framework is implemented using OSQP solver[77] and formulated as follows:

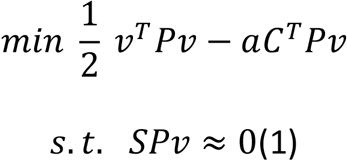

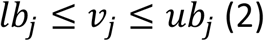

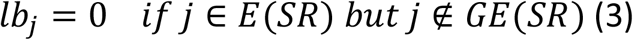

*C* is a vector of zeros with ones at each cell group’s designated biomass reaction position. *a* is set to 10000. Constraint (3) is the growth medium constraints, and we define a growth medium

*G*, and *GE(SR*) as shared exchange reactions relevant to *G. j* stands for the *jth* reaction.

### 4.5 Simulation

#### 4.5.1 Simulation of dropout data

To estimate the effect of data sparsity on model performance, we plan to employ a similar strategy described in splatter to simulate dropout on 11 NCI-60 cell lines we used[78]. First, we used a logistic function *f(x*) to calculate the dropout probability of every gene count.

Subsequently, a Bernoulli distribution with dropout probability vector as input parameter will then be utilized to randomly replace the original count matrix with 0.

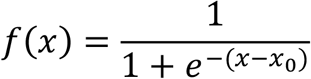

Where *x* is the log2 normalized gene count and *x*_0_ is the midpoint value. We increase the midpoint value sequentially to increase the proportion of dropout rates. The vector of midpoint values in the simulation is 0.1, 0.5, 1, 1.5, 2, 2.5, 3, 3. 5, 4, 4.5, 5. METAFlux will be applied for each simulated dataset and results obtained from simulated data will be compared with original data.

#### 4.5.2 Simulation of metabolic fluxes from random mediums

To estimate the medium effect on model performance, we will compare the results obtained from our biological meaningful medium with same-sized random mediums. The assumed medium contains 44 metabolites, so we randomly select 44 metabolites from the total 1648 exchange metabolites. We then allow our model to uptake or excrete those 44 metabolites without rate restriction while only allowing the rest of 1604 metabolites to excrete with no rate restriction. We repeat this process *N* = 500 times. For each simulation *i*, we obtain the overall Spearman correlation *ρ*_*i*_ and directionality accuracy *acc*_*i*_. To calculate the p-value, we simply count the number of measurements greater than our original biological meaningful statistics *ρ*_*o*_ and *acc*_*o*_ and divide this number by 500.

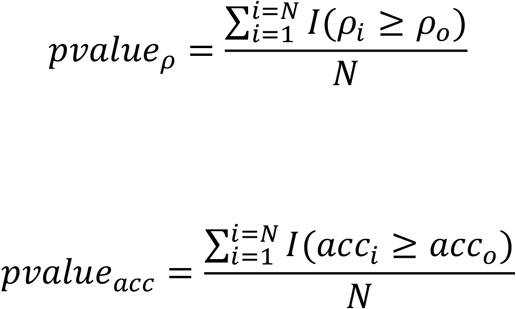

Where *I(*.) Is an indicator function, equaling 1 if the condition in parenthesis is true, 0 otherwise.

## 5 Additional Information

### 5.1 Ethics approval and consent to participate

Not applicable in this study.

### 5.2 Consent for publication

Not applicable in this study.

### 5.3 Data and materials availability

All datasets used in our study are publicly available. Human-GEM model was accessed from (https://github.com/SysBioChalmers/Human-GEM). We retrieved the medium composition and flux profiling data for 11 NCI-60 cell lines under the original manuscript [55]. ecGEM flux prediction for 11 NCI-60 cell lines was be obtained at https://zenodo.org/record/3583004#.YhQJdZPMJqs NCI-60 cell lines TPM RNA-seq data was obtained from https://depmap.org/portal/download/.The TCGA pan-cancer RNA-seq TPM data was downloaded from UCSC Xena data hubs(https://xenabrowser.net/). The proliferation score data can be found in the original publication[68]. CAR-NK single-cell RNA-seq data is available through NCBI Gene Expression Omnibus (GSE190976). Patient Lung cancer bulk sorted RNA-seq data can be downloaded from the NCBI Gene Expression Omnibus (GSE111907).

### 5.4 Code Availability

Code will be available in Github.

### 5.5 Competing interests

The authors declare that they have no competing interests.

### 5.6 Funding

This project has been made possible in part by grant RP180248 to KC from Cancer Prevention & Research Institute of Texas, grant U01CA247760 to KC and the Cancer Center Support Grant P30 CA016672 to PP from National Cancer Institute, and Human Cell Atlas Seed Network Grant CZF2019-02425 and CZF2019-002432 to KC from the Chan Zuckerberg Initiative DAF, an advised fund of Silicon Valley Community Foundation. This project was also partially supported by the MD Anderson Moonshot programs.

## 5.7 Acknowledgments

## Supplementary Data

### Supplementary tables

**Supplementary Table S1.** Nutrients identified in competing and cooperation mechanisms for patient lung cancer samples.

| Nutrients (Competing) | Nutrients(cooperating) |
| --- | --- |
| *Glucose, Isoleucine, Lysine, Methionine, Threonine, Tryptophan, O2, Glutamine, Tyrosine, Proline, Aspartate, Fe3+ Ornithine, Fructose, Folate, 2-hydroxybutyrate, Citrate, Carnitine, Hypoxanthine, Cystine* | *Histidine, Leucine, Phenylalanine, Valine, H2O, Asparagine, Cysteine, Arginine, Serine, Glutamate, PO43-, SO42-, Na+, K+, Ca2+, Glycerol, Acetate, Biotin, Galactose, Riboflavin, Pyridoxine, Cl-, Malonate, Citrulline, Acetone, Creatine, Formate, Betaine, Myo-Inositol, Niacinamide, Pyridoxal, Succinate, Urea* |

### Supplementary figures

**Supplementary Fig. S1.**
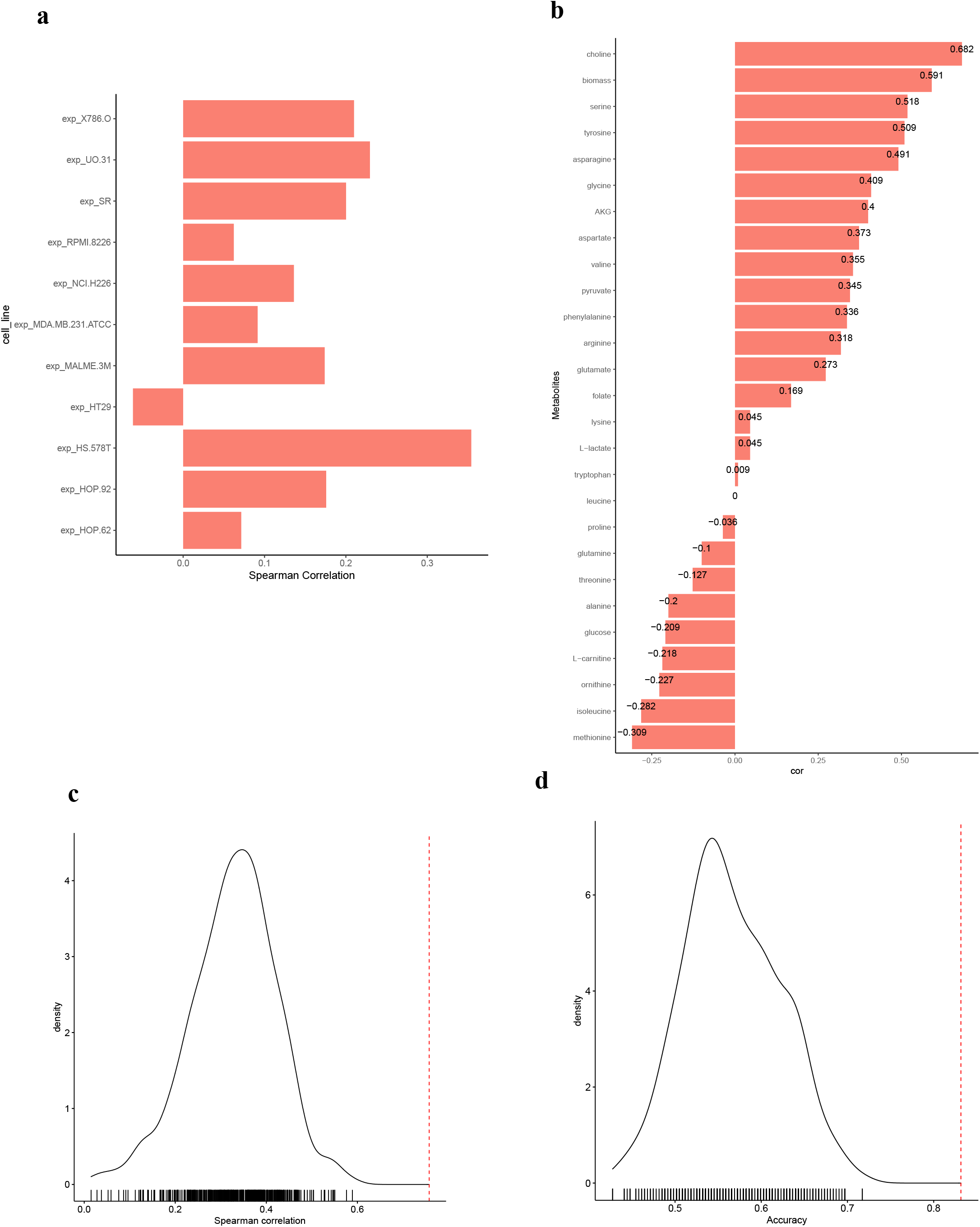
Simulation results of random medium and constraint-free medium predicted fluxes. **(a)** Spearman correlation bar plot across each cell line for METAFlux model without any medium constraints. The spearman correlations between predicted fluxes and experimental fluxes were calculated for 11 cell lines. **(b)** Spearman correlation bar plot across each metabolite for METAFlux model without any medium constraints. The spearman correlations between predicted fluxes and experimental fluxes were calculated for 26 metabolites and one biomass reaction. **(c)** Distribution of correlation between predicted fluxes with experimental fluxes over 500 random mediums. The red dotted line indicates the results of the original biological meaningful medium. **(d)** Distribution of direction accuracy over 500 random mediums. The red dotted line indicates the result of the original biological meaningful medium.

**Supplementary Fig. S2.**
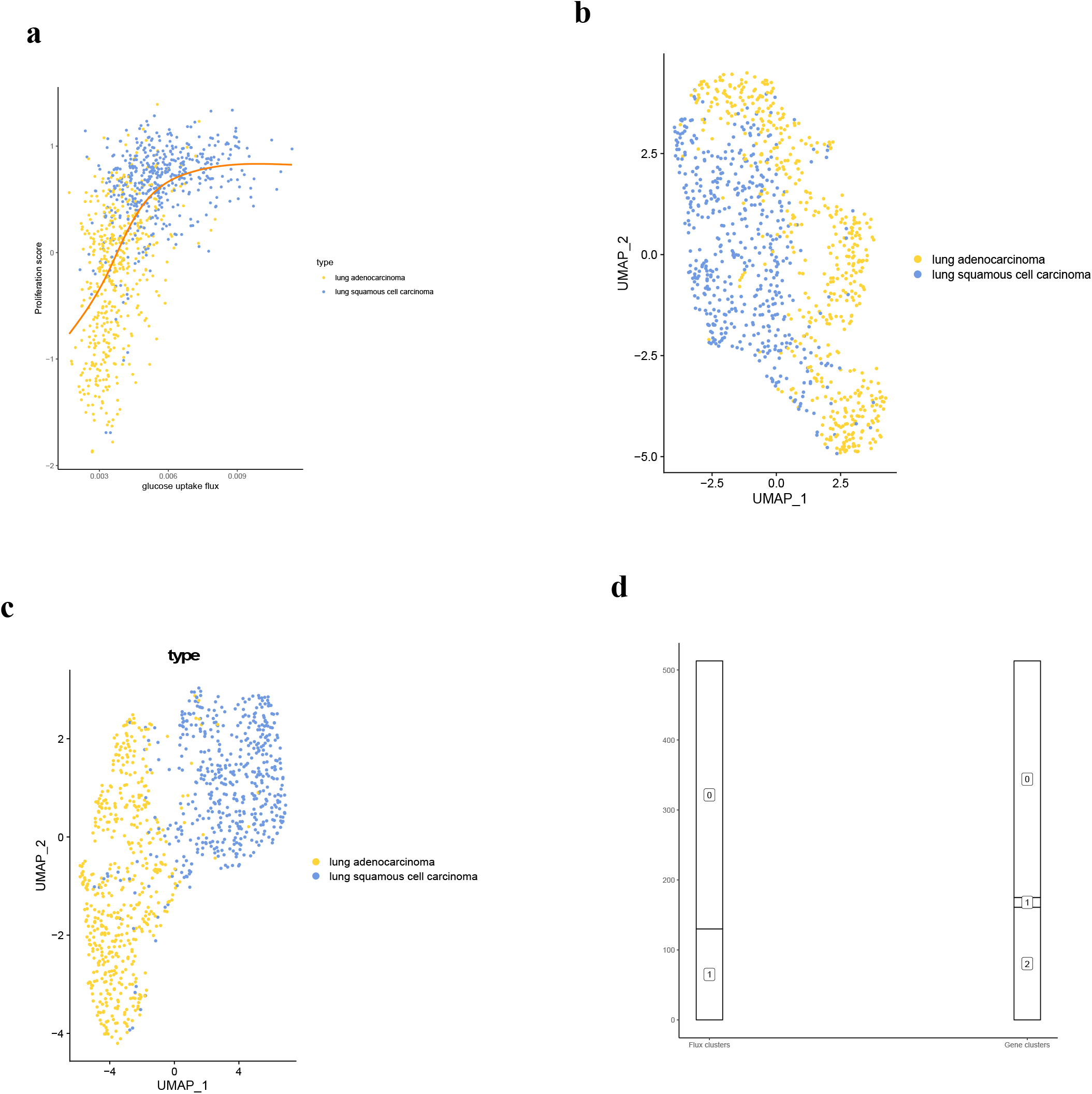
Clustering and UMAP of TCGA LUAD and LUSC. **(a)** Scatterplot of glucose uptake flux with proliferation score. Each dot was color-coded by lung cancer type. The trend curve was generated using loess smoothing. **(b)** UMAP of TCGA LUAD and LUSC samples using predicted metabolic fluxes. Each dot represents one sample, and it is color-coded by lung cancer type. **(c)** UMAP of TCGA LUAD and LUSC samples using metabolic gene expression. Each dot represents one sample, and it is color-coded by lung cancer type. **(d)** A river plot showing the clustering membership for flux-based clustering and gene-based clustering.

**Supplementary Fig. S3.**
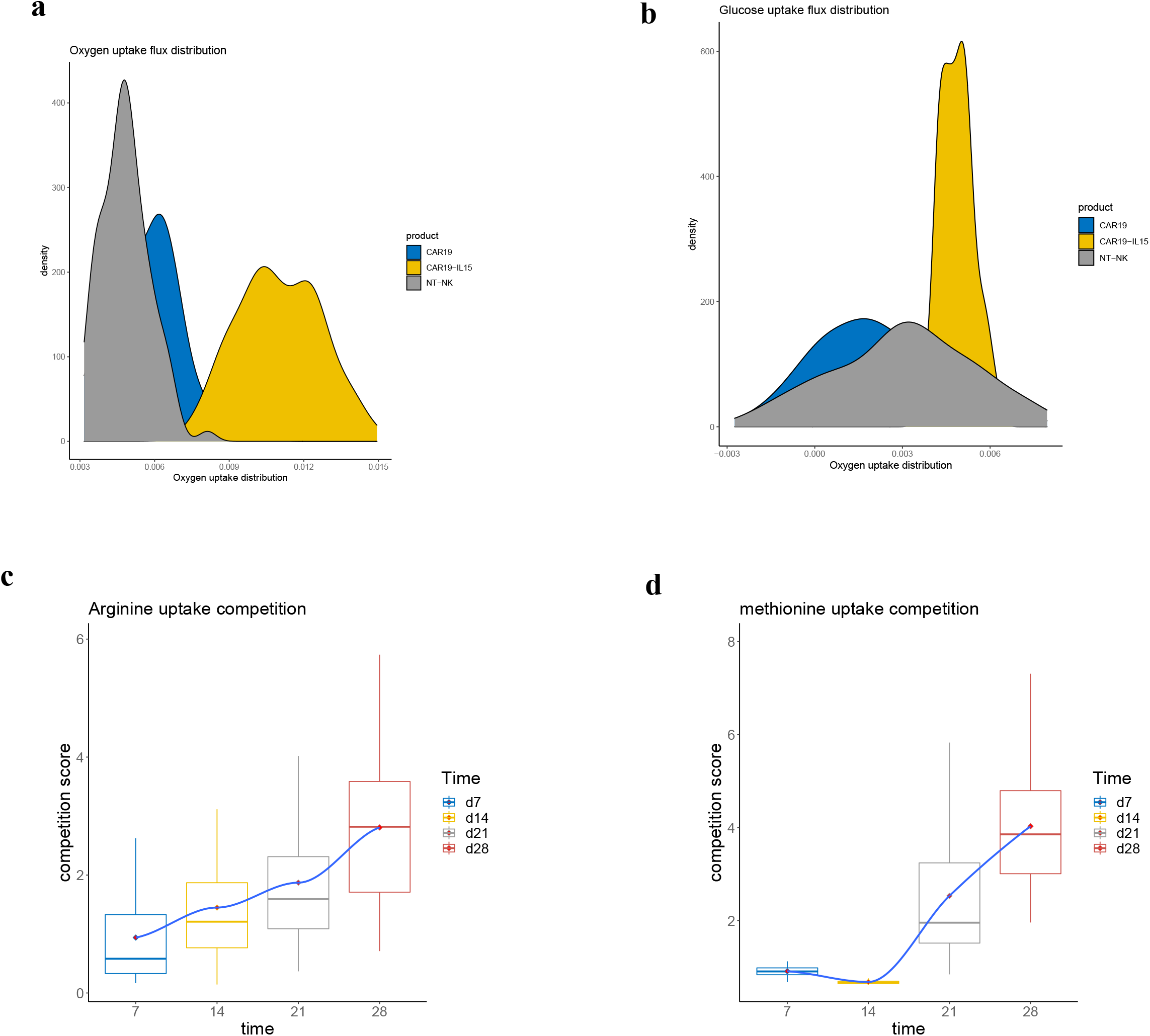
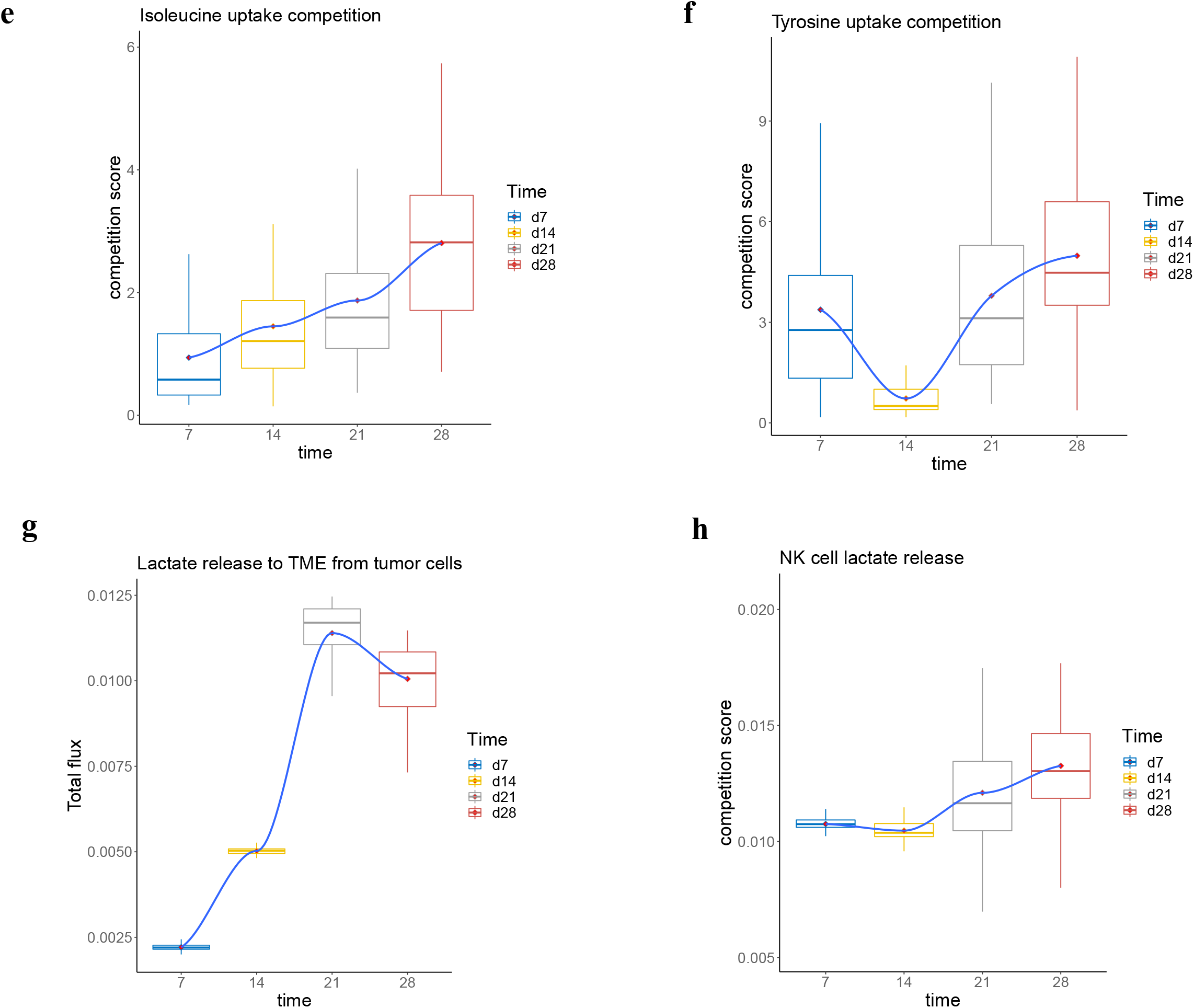
Nutrients competition and lactate release profile in CAR-NK datasets. **(a-b)** Distribution of glucose and oxygen fluxes for three products. **(c-f)** Competition score of **(c)** arginine, **(d)** methionine, **(e)** isoleucine, and **(f)** tyrosine for average cancer cell against average NK cell in TME from day 7 to day 28. **(g)** The trend plot of total tumor lactate release to TME from day 7 to day 28. **(h)** The trend plot of lactate release by average NK cell over time.

**Supplementary Fig. S4.**
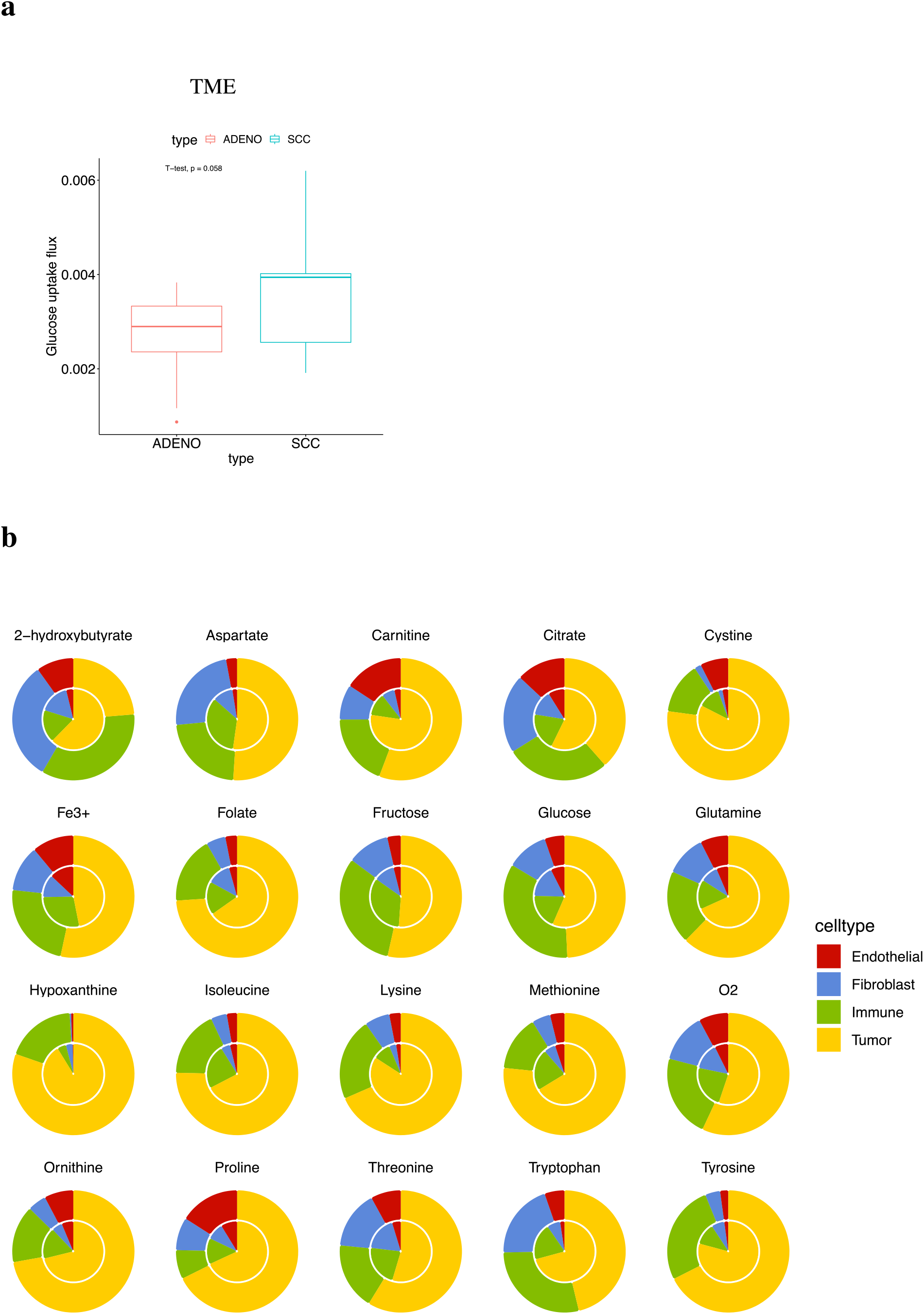
METAFlux results on patient lung cancer datasets. **(a)** The overall glucose uptake comparison for ADENO and SCC. **(b)** Donut pie charts of total cell type specific nutrient uptake percentage for ADENO and SCC. The outer circle shows the LUSC nutrient uptake, and the inner circle shows ADENO nutrient uptake percentage. To calculate the percentage of specific nutrient uptake in a particular cell type, we divided the total uptake of that nutrient from each cell type by the total TME uptake of that nutrient. The total cell type uptake was calculated using per cell average flux multiplied by cell type proportion. Total TME nutrient uptake was calculated by summing the total uptake of that nutrient from all cell types.

## Notes

### Competing Interest Statement

The authors have declared no competing interest.

